# Differential risk of incident cancer in patients with heart failure: A nationwide population-based cohort study

**DOI:** 10.1101/2020.01.10.901454

**Authors:** Soongu Kwak, Soonil Kwon, Seo-Young Lee, Seokhun Yang, Hyun-Jung Lee, Heesun Lee, Jun-Bean Park, Kyungdo Han, Yong-Jin Kim, Hyung-Kwan Kim

**Affiliations:** Department of Internal medicine, Seoul National University Hospital, Seoul, Korea; Healthcare System Gangnam Center, Seoul National University Hospital, Seoul, Korea; Department of Biostatistics, The Catholic University of Korea, Seoul, Korea

**Keywords:** Heart failure, Cancer, Incidence

## Abstract

**Background:** Heart failure (HF) and cancer are currently two leading causes of mortality, and sometimes coexist. However, the relationship between them is not completely elucidated. We aimed to investigate whether patients with HF are predisposed to cancer development using the large Korean National Health Insurance claims database.

**Methods and findings:** This study included 128,441 HF patients without a history of cancer and 642,205 age- and sex-matched individuals with no history of cancer and HF between 1 January 2010 and 31 December 2015. During a median follow-up of 4.06 years, 11,808 patients from the HF group and 40,805 participants from the control were newly diagnosed with cancer (cumulative incidence, 9.2% vs. 6.4%, *p*<0.0001). Patients with HF presented a higher risk for cancer development compared to controls in multivariable Cox analysis (hazard ratio [HR] 1.64, 95% confidence interval [CI] 1.61 - 1.68). The increased risk was consistent for all site-specific cancers. To minimize potential surveillance bias, additional analysis was performed by eliminating participants who developed cancer within the initial 2 years of HF diagnosis (i.e. 2-year lag analysis). In the 2-year lag analysis, the higher risk of overall cancer remained significant in patients with HF (HR 1.09, 95% CI 1.05 - 1.13), although the association was weaker. Among the site-specific cancers, three types of cancer (lung, liver/biliary/pancreas, and hematologic malignancy) were consistently at higher risk in patients with HF.

**Conclusions:** Cancer incidence is higher in patients with HF than in the general population. Active surveillance of coexisting malignancy needs to be considered in these patients.

## INTRODUCTION

Heart failure (HF) is one of the leading causes of mortality in developed countries [1]. Recent advances in the contemporary management of HF have improved survival [2]. Hence, the relative contributions of non-cardiovascular causes to mortality cannot be ignored in patients with HF [3,4].

Cancer is a major cause of mortality in the general population, and its incidence increases steadily with age. Due to the extended life expectancy of patients with HF, a new diagnosis of cancer in patients with established HF is not infrequent, and cancer is now one of the most common causes of non-cardiovascular mortality, accounting for up to 10% of the reported causes of death [3,4]. However, whether HF itself predisposes to an increased risk of malignancy has been rarely discussed compared to the development of HF due to cardiotoxicity induced by cancer chemotherapy [5].

Recently, four large cohort studies reported an increased incidence of cancer in patients with HF [6-9], whereas this association was dispelled in a study exclusively involving male physicians [10]. Several limitations such as the small sample size of the cohort [8], lack of appropriate risk adjustment [6,9], and short follow-up period [6,8], as well as the specific selection of the participants [10], may be the reasons why previous studies reported discrepant results. Moreover, the possibility of a surveillance bias, which may act as a major confounder within the studies, cannot be excluded, since active follow-up with frequent and regular hospital visit might provide a better opportunity for earlier and easier detection of cancer. Thus, the relationship between these two severe diseases is still in question, and more compelling evidence is warranted in a larger study population.

Thus, we aimed to evaluate the association between HF and cancer using data from the Korean National Health Insurance Service (NHIS) claims database.

## METHODS

### Data source and study population

We conducted a nationwide population-based cohort study using data from the Korean NHIS claims database. Korean NHIS system is a mandatory universal health insurance program managed by the Korean government since 1989 and offers comprehensive medical care to 97% of the Koreans [11]. The remaining 3% of Koreans with evidence of low income are covered by the Medical Aid Program, whose information has been incorporated into a single database since 2006. The NHIS database includes detailed information of an individual, including demographic characteristics, health behavior, diagnosis, prescription, surgery or procedures received, health care utilization (i.e. hospitalization) [11]. This well-constructed database was used in many previously published studies, and its validity as a reliable data source has been established [12-14].

Among the NHIS representative sample cohort between 2010 and 2015, the data of participants newly diagnosed with HF and aged ≥20 years were collected. HF was defined by claims for diagnostic codes (*International Classification of Disease, Tenth Revision, Clinical Modification; ICD-10-CM*) (I50) with at least one hospital admission attributed to HF diagnosis. Patients who had a history of cancer defined by *ICD-10-CM* codes (C00 to C97) at the time of enrolment were systematically excluded. For comparison, an age- and sex-matched control group comprising individuals who had no history of either HF or cancer was randomly selected (1:5 ratio). The flow diagram of patient selection is presented in **Fig 1.**

**Fig 1.**
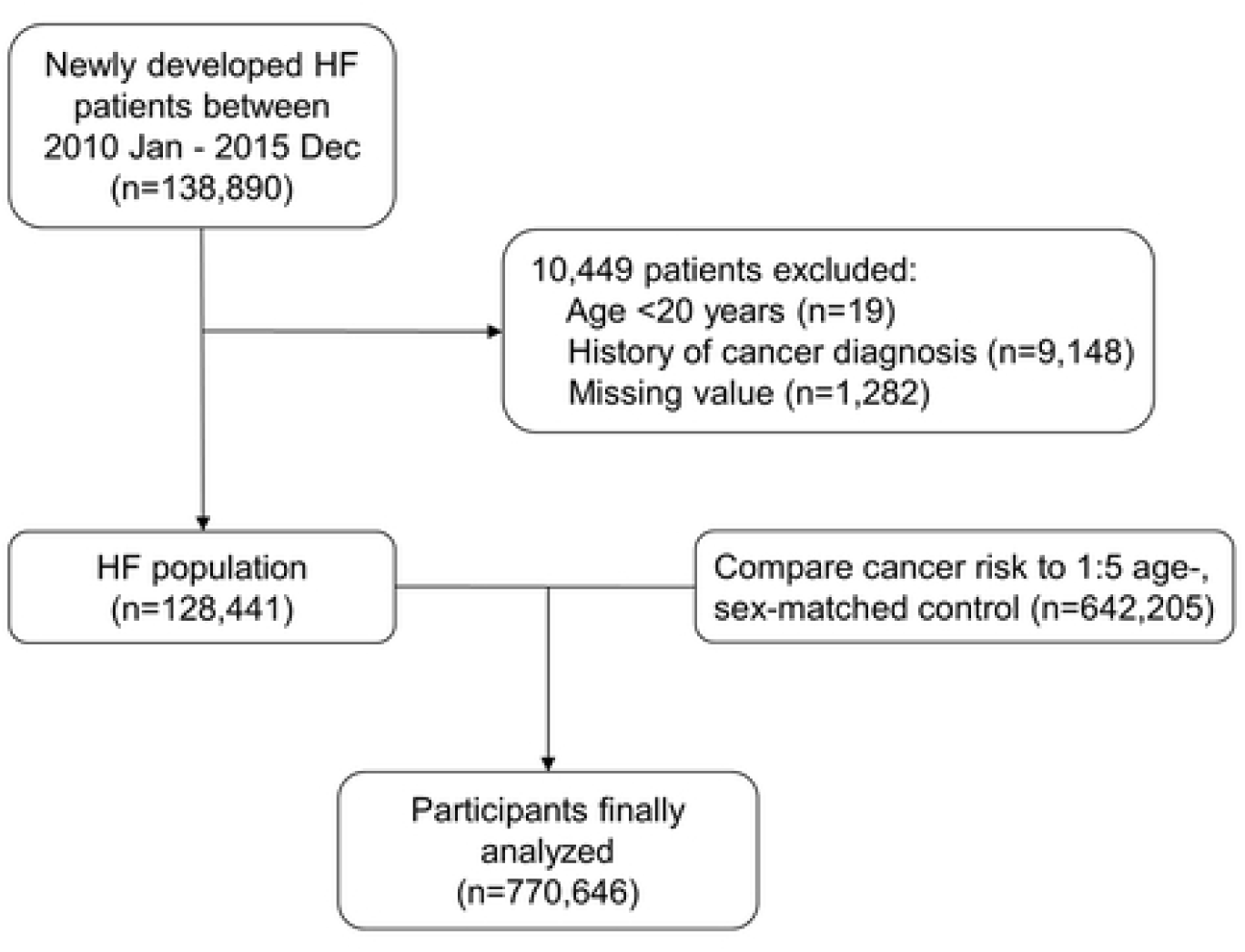
Flow chart for the selection of the study population. Flow chart depicting the process of patient selection included in the study. HF denotes heart failure.

The study protocol conformed to the ethical guidelines of the Declaration of Helsinki and was approved by the Institutional Review Board of our institution (Seoul National University Hospital, Seoul, Korea; IRB number, E-1806-018-949). As anonymized and unidentified information was used for the analysis, the need for informed consent was waived by the same ethic committee (Seoul National University Hospital).

### Diagnostic validity of cancer

In Korea, all cancers fall under the category of *Rare Intractable Diseases*; all patients in this category are designated as special medical aid beneficiaries with the expanding benefit of the NHIS. Since 2006, the government has introduced an initiative covering 90% of all medical expenses claimed by these patients. Therefore, the diagnosis of cancer is strictly determined and monitored by a thorough verification with clinical, imaging, and pathological evidence, and rigorous reviews by medical experts and health insurance professionals, according to an act established by the Ministry of Health and Welfare [13,14]. Therefore, data for cancer used in this study can be considered validated and reliable.

### Outcome measures

Participants with HF were followed up from the date of first HF diagnosis to the date of cancer diagnosis, or to the end of the study period (31 December 2017), whichever came first. For the controls, the follow-up time was from the date of assigned national health examination to the date of cancer diagnosis, or to the end of the study period (31 December 2017), whichever came first. Participants who died before cancer diagnosis during follow up were censored. Development of cancer was confirmed by both the new assignment of ICD-10-CM codes of cancer, and at the same period, new registration of the patient to the NHIS enhanced benefits coverage registry by the cancer diagnosis. The incidence of overall and each site-specific cancer was investigated. Site-specific cancers include gastrointestinal (GI) cancer (esophagus, stomach, colorectal), liver/biliary/pancreas cancer (liver, biliary, pancreas), lung cancer, prostate cancer, hematologic cancer (leukemia, lymphoma, multiple myeloma), genitourinary cancer (renal, bladder), thyroid cancer, breast cancer, female reproductive cancer (cervical, ovarian, uterine), head and neck cancer (oral, laryngeal), and skin cancer. ICD-10-CM codes of cancers used in this study are summarized in **S1 Table**.

### Statistical analysis

Categorical variables (frequencies and percentages) were compared using the χ^2^ test, and continuous variables (mean ± standard deviation or median with interquartile range) were analyzed by the Student’s *t-*test or Wilcoxon’s rank sum test for independent samples between the HF group and the control group. The incidence rates of cancer were calculated per 1,000 person-years. The cumulative incidence of cancer was plotted and compared between the HF group and the control group by the log-rank test. Cox proportional hazard regression analysis was performed to evaluate the association between HF and cancer development. The multivariable Cox models were adjusted for age, sex, income, diabetes mellitus, smoking, alcohol consumption, and body mass index. The adjusted hazard ratio (HR) was also calculated for the pre-specified subgroups (according to age, and the status of income, smoking, and drinking). The risk of cancer development in each site-specific cancer was expressed as HR with the corresponding 95% confidence interval (CI) in univariable and multivariable analyses.

To avoid potential surveillance bias, a sensitivity analysis was additionally performed by eliminating patients who were newly diagnosed with cancer within the initial 2 years of HF diagnosis, as well as those followed for less than 2 years (i.e. 2-year lag analysis). The same statistical analyses were repeated in this 2-year lag cohort. SAS software version 9.4 (SAS, Cary, NC, USA) was used in all statistical analyses, and *p* values <0.05 were considered statistically significant.

## RESULTS

### Study population

In the present cohort (n = 770,646; mean age, 67.1 years; men, 51.9%), patients with HF (n=128,441) were compared with age- and sex-matched controls (n = 642,205). **Table 1** summarizes the baseline characteristics of the study population. Briefly, patients with HF were more likely to be obese, to have a smoking history, and comorbidities such as diabetes mellitus, hypertension, and dyslipidemia (all *p* <0.0001), whereas alcohol consumption was more prevalent in the controls (*p* <0.0001).

**Table 1.** Baseline characteristics of the study population.

|  | HF<br>(n=128,441) | Control<br>(n=642,205) | *p* value |
| --- | --- | --- | --- |
| Male, n (%) | 66,687 (51.9) | 333,435 (51.9) | >0.99 |
| Age, years | 67.1±12.4 | 67.1±12.4 | >0.99 |
| 20-34 | 1,483 (1.1) | 7,415 (1.1) |  |
| 35-49 | 10,160 (7.9) | 50,800 (7.9) |  |
| 50-64 | 37,617 (29.3) | 188,085 (29.3) |  |
| ≥65 | 79,181 (61.7) | 395,905 (61.7) |  |
| Body mass index, kg/m<sup>2</sup> | 24.1 ± 3.6 | 23.8 ± 3.1 | <0.0001 |
| ≥25 | 49,708 (38.7) | 219,991 (34.2) | <0.0001 |
| Smoking, n (%) |  |  | <0.0001 |
| Never-smoker | 81,616 (63.5) | 424,483 (66.1) |  |
| Ex-smoker | 25,627 (19.9) | 119,582 (18.6) |  |
| Current-smoker | 21,198 (16.5) | 98,140 (15.2) |  |
| Alcohol, n (%) |  |  | <0.0001 |
| Never | 97,226 (75.7) | 435,031 (67.7) |  |
| Mild to moderate | 26,062 (20.2) | 177,241 (27.6) |  |
| Heavy* | 5,153 (4.0) | 29,933 (4.6) |  |
| Low income<sup>†</sup>, n (%) | 30,550 (23.7) | 131,267 (20.4) | <0.0001 |
| Diabetes mellitus, n (%) | 41,680 (32.4) | 124,574 (19.4) | <0.0001 |
| Hypertension, n (%) | 101,461 (78.9) | 353,761 (55.0) | <0.0001 |
| Dyslipidemia, n (%) | 70,959 (55.2) | 222,156 (34.5) | <0.0001 |

\*Heavy drinking status was defined by alcohol consumption ≥30 grams/day.
<sup>†</sup>Low income is defined as household income ≤30% of the median.
HF denotes heart failure.

### Incidence and cumulative incidence of cancer

The incidence of all cancers was higher in the HF group compared to that of the control group. During a median follow-up of 4.06 years (interquartile range, 2.75 – 5.76 years), 11,808 participants from the HF group and 40,805 participants from the control group were newly diagnosed with cancer (9.2% vs. 6.4%), corresponding to an incidence rate of 24.2 and 14.6 per 1,000 person-year, respectively (**Table 2**). The cumulative incidence of cancer in the HF group was higher than that of the control group (*p* <0.0001) (**Fig 2A**). The higher cancer incidence among the HF group was consistently observed for all site-specific cancers (**Table 2).** The cumulative incidence of the four most common site-specific cancers is shown in **Fig 3A.** Of note, the incidence rate of overall cancer was markedly increased in the HF group for the first 2 years of HF diagnosis (**Fig 2A**). The same trend was consistently observed for all major site-specific cancers (**Fig 3A**).

**Table 2.** Comparison of the incidence of cancer between the HF and the control group.

| Outcomes | No lag |  |  |  |  | 2-year lag |  |  |  |  |
| --- | --- | --- | --- | --- | --- | --- | --- | --- | --- | --- |
|  | HF (n=128,441) |  | Control (n=642,205) |  | *p* value | HF (n=101,924) |  | Control (n=578,266) |  | *p* value |
|  | Event | IR | Event | IR |  | Event | IR | Event | IR |  |
| Overall cancer | 11,808 (9.2) | 24.2 | 40,805 (6.4) | 14.6 | <0.0001 | 3,857 (3.8) | 8.4 | 22,127 (3.8) | 8.1 | 0.0001 |
| Site-specific cancers |  |  |  |  |  |  |  |  |  |  |
| Gastrointestinal | 4,236 (3.3) | 8.4 | 16,009 (2.5) | 5.6 | <0.0001 | 1,236 (1.2) | 2.6 | 8,004 (1.4) | 2.9 | 0.0615 |
| Liver/Biliary/Pancreas | 2,762 (2.2) | 5.4 | 8,460 (1.3) | 2.9 | <0.0001 | 859 (0.8) | 1.8 | 4,560 (0.8) | 1.6 | 0.0010 |
| Lung | 2,584 (2.0) | 5.0 | 6,725 (1.0) | 2.3 | <0.0001 | 781 (0.8) | 1.6 | 3,760 (0.7) | 1.3 | <0.0001 |
| Prostate* | 1,159 (1.7) | 4.5 | 4,986 (1.5) | 3.4 | <0.0001 | 420 (0.8) | 1.8 | 2,838 (1.0) | 2.0 | 0.9534 |
| Hematology | 924 (0.7) | 1.8 | 1,895 (0.3) | 0.6 | <0.0001 | 210 (0.2) | 0.4 | 1,111 (0.2) | 0.4 | 0.0033 |
| Genitourinary | 719 (0.6) | 1.4 | 2,628 (0.4) | 0.9 | <0.0001 | 250 (0.2) | 0.5 | 1,498 (0.2) | 0.5 | 0.2958 |
| Thyroid | 538 (0.4) | 1.0 | 2,135 (0.3) | 0.7 | <0.0001 | 149 (0.1) | 0.3 | 960 (0.2) | 0.3 | 0.4239 |
| Breast<sup>†</sup> | 386 (0.6) | 1.5 | 1,432 (0.5) | 1.0 | <0.0001 | 147 (0.3) | 0.6 | 721 (0.3) | 0.5 | 0.0484 |
| Female reproductive<sup>†</sup> | 354 (0.6) | 1.3 | 985 (0.3) | 0.7 | <0.0001 | 101 (0.2) | 0.4 | 496 (0.2) | 0.3 | 0.2213 |
| Head and neck | 248 (0.2) | 0.4 | 891 (0.1) | 0.3 | <0.0001 | 87 (0.09) | 0.1 | 476 (0.08) | 0.1 | 0.3684 |
| Skin | 49 (0.04) | 0.09 | 188 (0.03) | 0.06 | 0.0155 | 18 (0.02) | 0.03 | 114 (0.02) | 0.04 | 0.8730 |

\*Prostate cancer is analyzed only in men (n=66,687 in HF and n=333,435 in control for ‘no lag’ cohort; n=51,456 in HF and n=297,780 in control for ‘2-year lag’ cohort).
<sup>†</sup>Female reproductive malignancies and breast cancer are analyzed only in women (n=61,754 in HF and n=308,770 in control for ‘no lag’ cohort; n=50,468 in HF and n=280,486 in control for ‘2-year lag’ cohort).
HF denotes heart failure
IR denotes incidence ratio (1,000 person-year)

**Fig 2.**
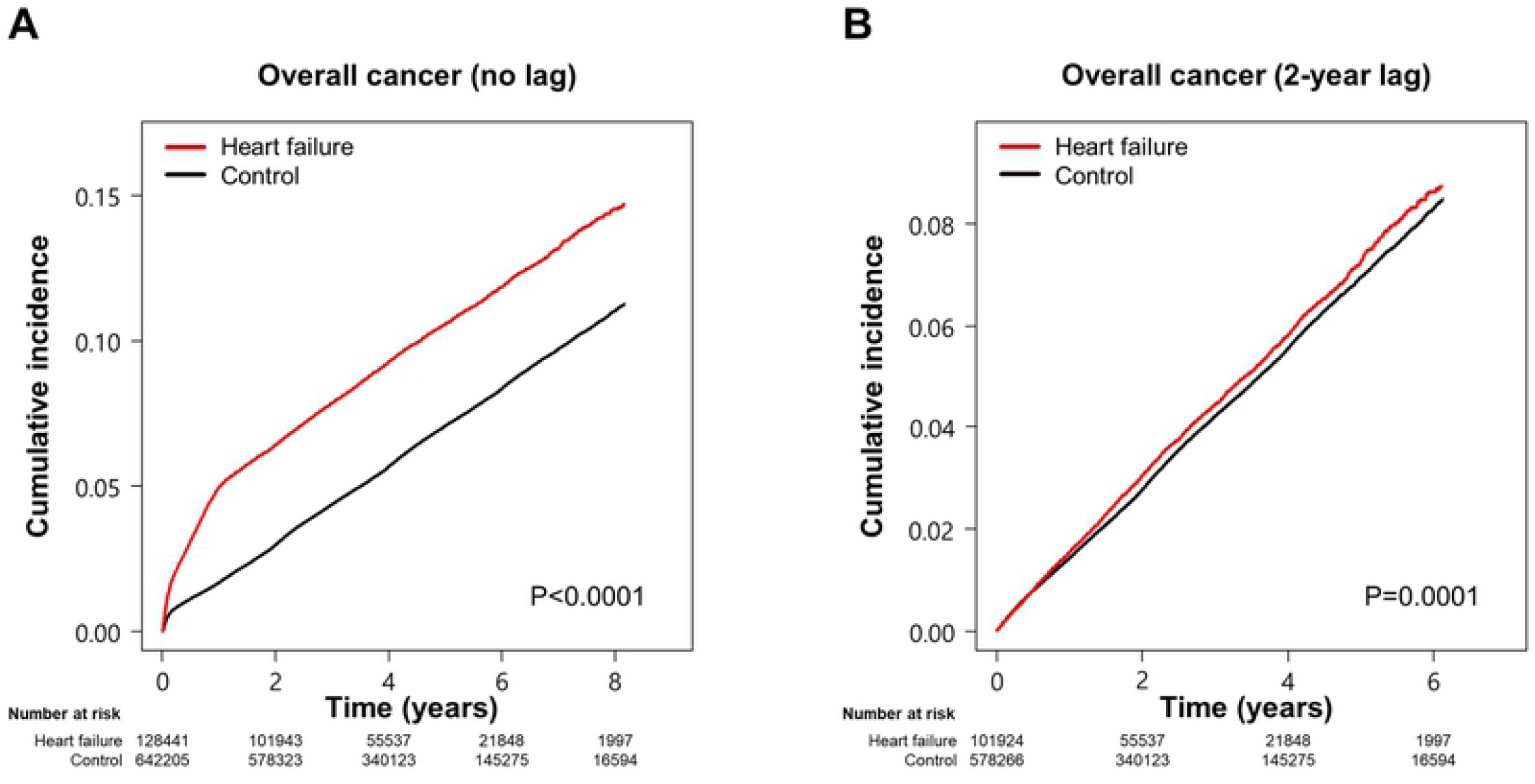
Cumulative incidence of overall cancer in the HF group and the control group. Kaplan-Meier curves of overall cancer incidence were compared between the HF group and the control group using the log-rank test. *A.* Overall cancer incidence after HF diagnosis; *B.* Overall cancer incidence after 2 year of HF diagnosis (2-year lag analysis). HF denotes heart failure.

**Fig 3.**
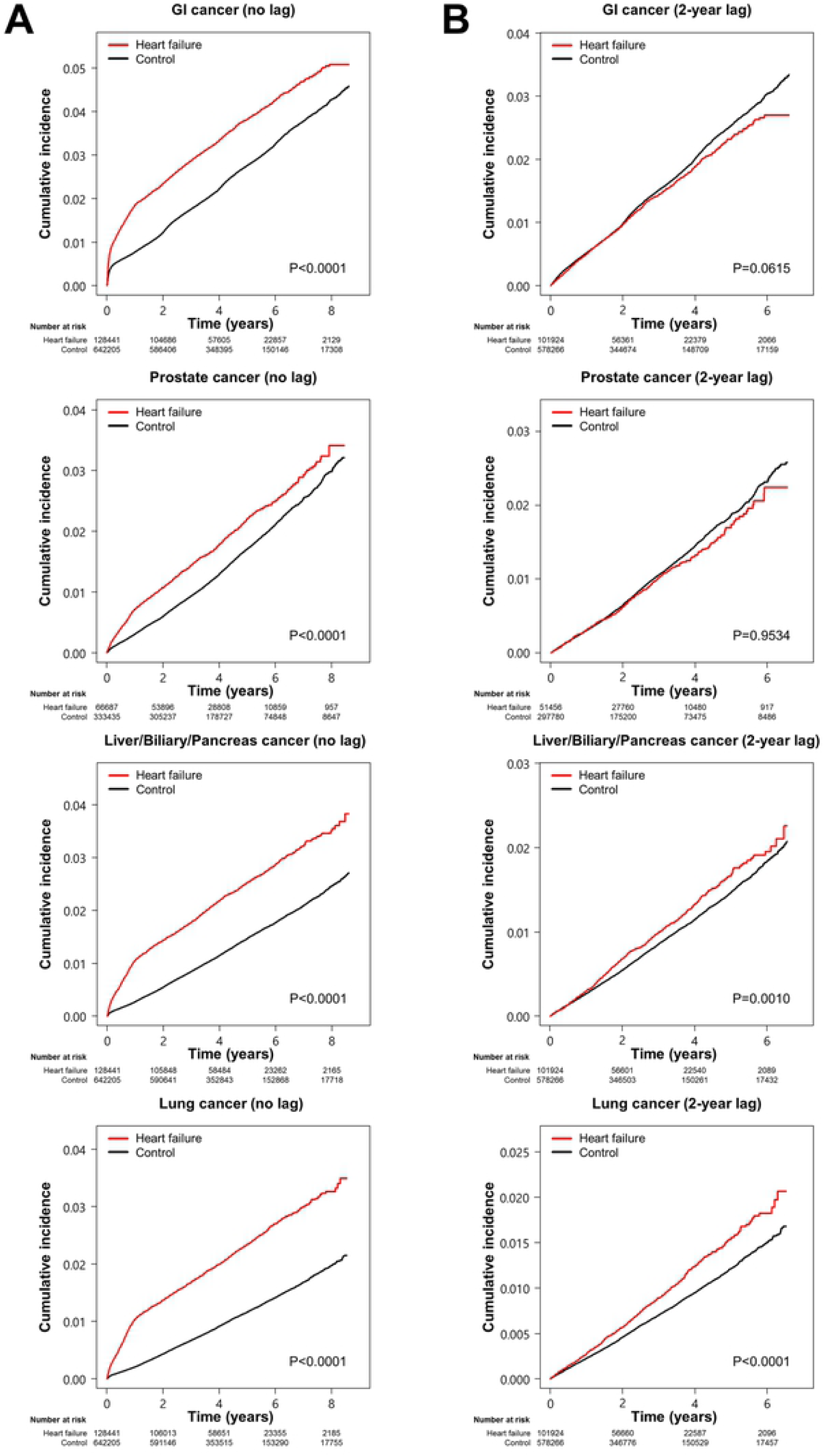
Cumulative incidence of site-specific cancers in the HF group and the control group. Kaplan-Meier curves for each site-specific cancer incidence were compared between the HF group and the control group using the log-rank test. *A.* Cumulative incidence of GI, prostate, liver/biliary/pancreas, and lung cancer in the no lag cohort. *B.* Cumulative incidence of GI, prostate, liver/biliary/pancreas, and lung cancer in the 2-year lag cohort. GI denotes gastrointestinal and HF, heart failure.

### Risk of cancer development

#### No lag analysis

During a median follow-up of 4.06 years (interquartile range, 2.75 – 5.76 years), the HF group showed an increased risk of overall cancer in the unadjusted analysis (HR 1.68, 95% CI 1.64-1.71; *p* <0.0001), and in multivariable-adjusted analysis (HR 1.64, 95% CI 1.61 - 1.68; *p* <0.0001) (**Table 3**). The risk of all site-specific cancers was consistently higher in the HF group in both the unadjusted and multivariable-adjusted analysis (**Table 3**), with the highest HRs noted for hematologic and lung malignancy. This association remained constant in the separate analysis by sex except for skin cancer (**S2 Table**). The adjusted risk of overall cancer was also significantly increased in all pre-specified subgroups (**S1 Fig**).

**Table 3.** Association of heart failure with cancer development in the ‘no lag’ and ‘2-year lag’ analysis.

| Outcomes | No lag |  |  |  | 2-year lag |  |  |  |
| --- | --- | --- | --- | --- | --- | --- | --- | --- |
|  | Unadjusted | *p* value | Adjusted* | *p* value | Unadjusted | *p* value | Adjusted* | *p* value |
| Overall cancer | 1.68 (1.64-1.71) | <0.0001 | 1.64 (1.61-1.68) | <0.0001 | 1.11 (1.08-1.15) | <0.0001 | 1.09 (1.05-1.13) | <0.0001 |
| Site-specific groups |  |  |  |  |  |  |  |  |
| Gastrointestinal | 1.51 (1.46-1.56) | <0.0001 | 1.49 (1.44-1.54) | <0.0001 | 0.99 (0.93-1.05) | 0.7839 | 0.97 (0.91-1.03) | 0.3794 |
| Liver/Biliary/Pancreas | 1.90 (1.82-1.98) | <0.0001 | 1.80 (1.72-1.88) | <0.0001 | 1.22 (1.13-1.31) | <0.0001 | 1.16 (1.08-1.25) | <0.0001 |
| Lung | 2.26 (2.16-2.37) | <0.0001 | 2.22 (2.12-2.32) | <0.0001 | 1.38 (1.28-1.49) | <0.0001 | 1.35 (1.25-1.46) | <0.0001 |
| Prostate<sup>†</sup> | 1.40 (1.32-1.50) | <0.0001 | 1.40 (1.31-1.49) | <0.0001 | 1.01 (0.91-1.12) | 0.8304 | 1.01 (0.91-1.12) | 0.8040 |
| Hematology | 2.79 (2.58-3.02) | <0.0001 | 2.77 (2.55-3.00) | <0.0001 | 1.22 (1.05-1.42) | 0.0065 | 1.20 (1.03-1.39) | 0.0159 |
| Genitourinary | 1.61 (1.48-1.75) | <0.0001 | 1.55 (1.43-1.69) | <0.0001 | 1.10 (0.96-1.26) | 0.1515 | 1.05 (0.92-1.20) | 0.4423 |
| Thyroid | 1.34 (1.22-1.48) | <0.0001 | 1.30 (1.18-1.43) | <0.0001 | 0.89 (0.75-1.06) | 0.1949 | 0.84 (0.70-1.00) | 0.0545 |
| Breast<sup>‡</sup> | 1.44 (1.28-1.61) | <0.0001 | 1.36 (1.21-1.52) | <0.0001 | 1.16 (0.97-1.39) | 0.0905 | 1.09 (0.91-1.31) | 0.3143 |
| Female reproductive<sup>‡</sup> | 1.95 (1.72-2.20) | <0.0001 | 1.90 (1.68-2.15) | <0.0001 | 1.19 (0.96-1.48) | 0.0979 | 1.14 (0.92-1.42) | 0.2164 |
| Head and neck | 1.61 (1.40-1.86) | <0.0001 | 1.62 (1.41-1.87) | <0.0001 | 1.18 (0.93-1.48) | 0.1556 | 1.19 (0.94-1.50) | 0.1354 |
| Skin | 1.52 (1.11-2.09) | 0.0085 | 1.53 (1.11-2.11) | 0.0081 | 1.01 (0.61-1.67) | 0.9496 | 1.00 (0.60-1.65) | 0.9835 |

Values are expressed as hazard ratios (95% confidence interval).
\*Adjusted by age, sex, income, diabetes mellitus, smoking, alcohol consumption and body mass index.
†Prostate cancer is analyzed only in men (n=66,687 in HF and n=333,435 in control for ‘no lag’ cohort; n=51,456 in HF and n=297,780 in control for ‘2-year lag’ cohort).
‡Female reproductive malignancies and breast cancer are analyzed only in women (n=61,754 in HF and n=308,770 in control for ‘no lag’ cohort; n=50,468 in HF and n=280,486 in control for ‘2-year lag’ cohort).
HF denotes heart failure.

#### 2-year lag analysis

To avoid the potential surveillance bias, the same analyses were repeated after eliminating patients with either cancer diagnosis established within 2 years of HF diagnosis or having a follow-up duration of less than 2 years. About 80% of patients remained in the HF group (n = 101,924), and 578,266 participants remained in the control group for comparison. Similar to the original cohort, the HF group in the 2-year lag cohort showed a higher prevalence of comorbidities, including obesity, smoking history, diabetes mellitus, hypertension, and dyslipidemia, except for alcohol consumption (**S3 Table).**

The mean follow-up period of the 2-year lag cohort was 4.48 years (interquartile range, 3.17 – 5.97 years). The number of cancer diagnoses and incidence rate per 1,000 person-year in the 2-year lag cohort are shown in **Table 2**. The cumulative incidence of overall cancers in the 2-year lag cohort was still significantly higher in the HF group (*p* =0.0001), although the gap between the two groups became smaller than that analyzed in the original cohort (**Fig 2B**). Cox proportional hazard analysis showed that the HF group demonstrated a significantly increased risk of overall cancers in both the unadjusted (HR 1.11, 95% CI 1.08 - 1.15; *p* <0.0001) and multivariable-adjusted analysis (HR 1.09, 95% CI 1.05 - 1.13; *p* <0.0001), with a smaller HR compared to that of no lag analysis (**Table 3**). The adjusted risks of all cancers in all pre-specified subgroups are illustrated in **S2 Fig**.

With regard to the site-specific cancers, the risk remained higher for liver/biliary/pancreas, lung, and hematologic malignancies, while the statistical difference was lost for the other site-specific cancers in this analyses (**Table 3**). The cumulative incidence of the four most common site-specific cancers in this 2-year lag analysis is shown in **Fig 3B**.

## DISCUSSION

In this study, we evaluated the relationship between HF and cancer development by analyzing data from a large Korean NHIS claims database. The current population-based cohort study has two main findings. Firstly, an abrupt increase of the new cancer diagnosis was observed in the first 2 years of HF diagnosis. Subsequently, the 2-year lag analysis (performed to minimize potential surveillance bias) proved a higher risk of overall cancer in the HF group compared to the control group, although the difference in the cancer incidence was much smaller. Secondly, the association of HF with the development of site-specific cancers was variable. While the ‘no lag’ analysis showed consistently increased risk of all site-specific cancers in both the unadjusted and multivariable-adjusted models, three subtypes of malignancies (liver/biliary/pancreas, lung, and hematologic malignancies) remained at higher risk in the ‘2-year lag’ unadjusted and multivariable-adjusted analyses.

A decade ago, HF and cancer were considered unrelated, with no influence on each other. Recent studies, however, suggested a shared pathophysiologic link between HF and cancer. For instance, chronic inflammation is a well-established mechanism of cancer development [15], which can also act as a crucial disease modifier in HF [16,17]. This is advocated by the elevated levels of pro-inflammatory cytokines that are also closely associated with the adverse outcomes in patients with HF [17-19]. This plausible hypothesis is further supported by a higher risk of cancer in other chronic inflammatory disorders [6,20,21]. More recently, Meijers *et al.* reported that precancerous lesions developed more frequently in HF-induced mice [22]. They found that several proteins, such as serpin A3 and A1, fibronectin, ceruloplasmin, and paraoxonase 1, were associated with enhanced tumor growth, independent of hemodynamic impairment, suggesting a potential mechanism of tumorigenesis induced by circulating factors produced by the failing heart [22]. In addition, they proved that elevated levels of inflammatory biomarkers in healthy participants had a predictive value for new-onset cancer independent of cancer-related risk factors, such as age, smoking status, and obesity [22]. Thus, the study provided evidence that HF is closely related to cancer development by enhanced inflammation.

An increased risk of cancer in HF patients was previously suggested in four cohort studies. In 2013, Hasin *et al.* reported a 68% higher risk of cancer development in a cohort including 596 patients with HF after adjusting for cancer-related risk factors, including obesity, smoking status, and comorbidities [7]. This finding was reproduced in a cohort of HF patients caused by myocardial infarction (n = 1,081) [8], in the Danish HF cohort (n = 9,307) [6] and in the Japanese population (n=5,238) [9]. In contrast, only one study refuted the association between HF and cancer development [10]. The study exclusively enrolled male population and used a patients-self report for HF and cancer diagnosis, and as such this issue is still under debate. Hence, additional, sizable studies with diagnostic validation and inclusion of both sexes are required to establish the association between these two grave diseases [23].

The current study presented an association between HF and cancer, with a HR of 1.64 for overall cancer and HRs in the range of 1.30 to 2.77 for site-specific cancer subtypes in the ‘no lag’ analysis. The most powerful advantage of our study compared to previous reports was the largest sample size of the cohort including 128,441 patients with HF and 642,205 community-based age- and sex-matched controls. In addition, a cancer diagnosis was verified by strict nationwide monitoring due to unique national insurance system and *Rare Intractable Disease* program of Korea, enabling accurate estimation of cancer development. More importantly, ‘2-year lag’ analysis was performed to minimize any possibility of co-existence of HF and hidden or subclinical malignancy, and the surveillance bias. In fact, during the first 2 years after HF diagnosis, the detection rate of cancer was substantially elevated compared to that of the general population, suggesting that surveillance bias may play a role due to the intensified medical evaluation. Nevertheless, the HF group still presented an increased cancer risk in the ‘2-year lag’ analysis, although the association was weaker, supporting the link between the two diseases.

Of interest, each site-specific cancer had different risks in the 2-year lag adjusted model; specifically, hematologic malignancy, lung cancer, and liver/biliary/pancreas cancer presented persistently elevated risk. Similar findings were demonstrated in the Danish HF cohort study [6]. This result implies that the impact of HF on cancer development may differ, and we can speculate that the mechanisms involved in the link between the two devastating disorders can be multifactorial.

Clinicians are apt to focus on cardiovascular consequences in patients with HF. However, with improved survival in HF and the aging societies, cancer has become non-negligible morbidity among the patients with HF [24], accounting for 10% of total deaths [3,4]. Outcomes of superimposed cancer in patients with HF are more dismal than in patients with HF alone [7], or cancer patients without HF [6,25]. Furthermore, the clinical decision can be modified in the presence of the concurrent malignancy; for example, the decision to implant a defibrillator or a left ventricular assist device can be rejected in patients diagnosed with cancer at an advanced stage [26]. Therefore, timely detection of de novo malignancy in patients with HF can be critical. Our findings demonstrated the higher cancer incidence in patients with HF than that of the general population, which implies that physicians are more encouraged to keep the possibility of coexisting malignancy in mind, and to perform active cancer screening in these population at risk. In particular, the surveillance could be targeted on the certain types of malignancies (i.e., lung cancer), given the differential risk of each site-specific cancer. Future studies are warranted to investigate whether surveillance of cancer screening can lead to an improvement in clinical outcomes in patients with HF.

This study is not without limitations. First, more detailed information of cancer-associated covariates, such as a family history of cancer and physical activity, was lacking. Medication data regarding cardiovascular drugs, such as angiotensin-converting enzyme inhibitors, angiotensin-receptor blockers, and beta-blockers, were also unavailable. Although a positive correlation between some HF medications, like angiotensin-receptor blockers, and cancer risk was previously reported [27,28], more recent publications refuted their associations [29-31]. Second, the estimation of left ventricular ejection fraction was not available. Although no substantial difference in the risk of cancer by left ventricular ejection fraction was reported, data on left ventricular systolic function would have strengthened our data [7,8,9]. Finally, the median follow-up of 4.06 years may not be long enough to adequately assess the impact of HF on cancer development. However, we believe that this limitation can be overcome by the large sample size of our cohort and by complete follow-up data obtained because of the unique medical reimbursement system of our country.

## CONCLUSIONS

This sizable, population-based cohort study found that patients with HF may carry a substantial risk of cancer development. In particular, the risk of liver/biliary/pancreas, lung, and hematologic malignancies was increased, even after excluding HF patients who developed cancer within 2 years after HF diagnosis in order to minimize potential surveillance bias. Therefore, from the clinical point of view, increased awareness and active surveillance of malignancy need to be considered in patients with HF.

## Acknowledgments

None

## Author Contributions

Conceptualization: Soongu Kwak and Hyung-Kwan Kim;

Data curation: Soongu Kwak, Soonil Kwon, Seo-Young Lee, Seokhun Yang, Hyun-Jung Lee,

Heesun Lee, Jun-Bean Park, and Hyung-Kwan Kim;

Formal analysis: Soongu Kwak, Kyungdo Han, and Hyung-Kwan Kim;

Funding acquisition: Hyung-Kwan Kim

Investigation: Soongu Kwak, Kyungdo Han, and Hyung-Kwan Kim;

Methodology: Kyungdo Han and Hyung-Kwan Kim;

Project Administration: Soongu Kwak and Hyung-Kwan Kim;

Resources: Kyungdo Han and Hyung-Kwan Kim;

Software: Kyungdo Han;

Supervision: Yong-Jin Kim and Hyung-Kwan Kim;

Validation: Kyungdo Han and Hyung-Kwan Kim;

Visualization: Soongu Kwak;

Writing – Original Draft Preparation: Soongu Kwak and Hyung-Kwan Kim;

Writing – Review & Editing: Soongu Kwak, Soonil Kwon, Seo-Young Lee, Seokhun Yang,

Hyun-Jung Lee, Heesun Lee, Jun-Bean Park, Yong-Jin Kim, and Hyung-Kwan Kim;

## Supporting information

**S1 Table. ICD-10 codes of overall and subtypes of cancers.** (DOCX)

**S2 Table. Association of heart failure with cancer development in a separate analysis by sex.** (DOCX)

**S3 Table. Baseline characteristics of the study population excluding patients with either cancer diagnosis within 2 years of HF diagnosis or having follow-up duration of less than 2 years.** (DOCX)

**S1 Fig. Association of cancer risk with HF in major subgroups (no lag cohort).** (DOCX)

**S2 Fig. Association of cancer risk with HF in major subgroups (2-year lag cohort).** (DOCX)

